# The loss-of-function of SOCS2 increases the inflammatory response to *Staphylococcus aureus* infection

**DOI:** 10.1101/2023.08.22.554270

**Authors:** Laurence Guzylack-Piriou, Blandine Gausseres, Christian Tasca, Chervin Hassel, Guillaume Tabouret, Gilles Foucras

## Abstract

The involvement of suppressor of cytokine signaling (SOCS)2 in anti-infective bacterial immunity remains largely undetermined compared to other members of the SOCS family. We developed a mouse model expressing the loss of function R96C SOCS2 point mutation to characterize the response of macrophages to *Staphylococcus aureus* and its TLR-ligand derivatives. The model resumes observations of gigantism done in Socs2-/- mice. Stimulation of bone-marrow-derived macrophages with various TLR-2 ligands showed upregulation of the pro-inflammatory cytokines IL-6 and TNF-α production only in cytokine-modulating environments that promote SOCS2 expression. Using this model, we showed that SOCS2 protein reduces STAT-5 phosphorylation in a short time frame upon TLR engagement. When SOCS2 is ablated, neutrophil and F4/80^int^ Ly6C^+^ inflammatory macrophage recruitment, as well as IFN-γ and IL-10 concentrations are significantly increased upon *S. aureus* peritoneal infection. By lowering the pro-inflammatory environment, SOCS2 favors better healing during a systemic infection caused by *S. aureus*.

## INTRODUCTION

In recent years, it has become increasingly evident that suppressor of cytokine signaling (SOCS) proteins have important roles in the maintenance of homeostasis and resolution of inflammatory processes. The two main functional domains of this protein family include the SH2 region and a C-terminal SOCS box. The SOCS box mediates the assembly of elongin B/C-cullin complexes to facilitate ubiquitination processes, leading to proteasomal degradation (Linossi & Nicholson, 2012). Most importantly, the SH2 domain interacts with its substrate by recognizing phosphorylated tyrosine residues and enhances substrate interaction through the N-terminal extended SH2-subdomain. As a member of the SOCS family, SOCS2 is a well-established negative regulator of growth hormone signaling via the JAK2/STAT-5 pathway (Li *et al*, 2022; Horvat & Medrano, 2001) and docks to the intracellular domains of related receptors or facilitates proteasome-dependent degradation of transcription factors (Vesterlund *et al*, 2011). The SH2 domain of SOCS2 can only bind to phosphorylated tyrosine residues on the target (Linossi *et al*, 2021). Indeed, the SOCS2-SH2 domain has been reported to directly interact with JAK2 (Kim *et al*, 2017). Thus, SOCSs proteins are likely to be involved in the differentiation of cells involved in innate and adaptive immunity, thus helping to shape the inflammatory response (Palmer & Restifo, 2009). Importantly, SOCS proteins appear to modulate CD4^+^ T-cell polarization (Knosp *et al*, 2011). This is exemplified by the regulation of the differentiation of Th2 cells by SOCS3 and SOCS2 (Knosp *et al*, 2011; Seki *et al*, 2003), whereas Th17 differentiation has been shown to be regulated by SOCS1 and SOCS3 (Chen *et al*, 2006; Tanaka *et al*, 2008). Moreover, Knosp et al. (Knosp *et al*, 2013) showed a dual role for SOCS2 in both Th2 and Foxp3^+^ iTreg generation.

Several studies using Socs2^−/−^ mice have highlighted the contribution of SOCS2 in regulating immune cell function in specific inflammatory or infectious contexts (Esper *et al*, 2012). SOCS2 deficiency induces hyper-responsiveness of dendritic cells (DCs) to microbial stimuli (Machado *et al*, 2006) and is related to an unbalanced inflammatory response during *Toxoplasma gondii*, *Trypanosoma cruzi*, and *Plasmodium berghei* infections (Brant *et al*, 2016). The action of the anti-inflammatory drug acetylsalicylic acid was shown to be dependent on SOCS2, confirming its role in both the immune response to infection and the regulation of inflammatory processes (Machado *et al*, 2006).

A recent study identified a mutation in the *Socs2* gene in ewes with an increased mammary inflammatory response (Rupp *et al*, 2015) and a higher prevalence of mammary infections by *Staphyloccoci*. This mutation changes the amino acid arginine at position 96, located in the SOCS2-SH2 domain, into cysteine (p.R96C), leading to disruption of the phospho-peptide binding pocket and the loss of binding to the ligand, further demonstrating the contribution of SH2:pTyr binding to the function of SOCS2 *in vivo* (1,15).

The role of the SOCS2 protein during bacterial infection has been little investigated. Few reports have noted their involvement in the regulation of Gram-positive bacteria-associated inflammation and that by Gram-negative bacteria, as well as lipopolysaccharide (LPS) signaling (Duncan *et al*, 2017). Unlike SOCS1 and SOCS3, the control of LPS signaling by SOCS2 is minimal. To promote TLR4 signaling, SOCS2 may target and mediate proteasome-dependent degradation of SOCS1 and SOCS3 (Tannahill *et al*, 2005; Hu *et al*, 2009). The expression of SOCS2 was shown to increase in macrophages infected with mycobacteria and SOCS2-deficient mice exhibit higher sensitivity to the inflammation induced by *Mycobacterium. bovis* (Carow & Rottenberg, 2014). Nonetheless, most studies have concluded that the activity of SOCS2 is limited and redundant, in contrast to the higher predisposition of SOCS2-deprived sheep to Staphylococcal infections.

The identification of the loss-of-function R96C mutation as a predisposing factor for mastitis induced by Gram-positive bacteria provides an opportunity to better understand the role of SOCS2 in the context of infection. We developed a genetically edited mouse model to express the SOCS2^R96C^ mutation to characterize the role of SOCS2 during the response to bacterial infection using *ex vivo* and *in vivo* experimental models. Here, we show that SOCS2 plays a more important role in the regulation of the response to infection and inflammatory processes than previously reported.

## RESULTS

### Introgression of the loss-of-function R96C-SOCS2 point mutation into the mouse genome resembles the *Socs2* knockout phenotype

We investigated the impact of the R96C mutation *in vivo* using CRISPR/Cas9 gene editing to generate a C57BL/6 mouse strain bearing the Arg96 to Cys mutation (SOCS2^KI^ mice). SOCS2^KI^ mice were indistinguishable from wildtype (WT) mice until weaning at three weeks of age but subsequently grew more rapidly and achieved higher body weight (Figure 1a). By six weeks of age, SOCS2^KI^ males and females weighed significantly more than their WT counterparts and adult mice were, on average, 40% heavier (Figure 1b). CT imaging of skeletons from 10-week-old males revealed that carcass weight was higher, indicating that muscle and bone may contribute significantly to the greater size of SOCS2-R96C mice. Consistent with this interpretation, the femur, tibia, radius, and ulna, as well as the skull in SOCS2^KI^ mice were all significantly longer than in WT controls (Figure 1c). We next characterized the immune-cell composition of various immune compartments, such as the spleen, LNs, and peritoneal cavity of SOCS2^KI^ mice relative to those of WT mice (Supplemental Figure 1). There were no significant differences in immune-cell composition in the spleen (Figure 1d), LNs (Figure 1e), or peritoneal cavity (Figure 1f). However, the total cell number of cells in the spleen was significantly lower in the SOCS2^KI^ mice when the calculation was established based on body weight (Supplemental Figure 2). Recent studies have highlighted the important role of macrophages during *S. aureus* infection (Pidwill *et al*, 2021). We first analyzed macrophage/DC subpopulations in the peritoneal cavity and spleens of SOCS2^KI^ mice relative to those of WT mice (Figure 1g, Supplemental Figure 3). At steady state, there were no significant differences in the various subpopulations of macrophages or DCs between the two genotypes.

**Figure 1.**
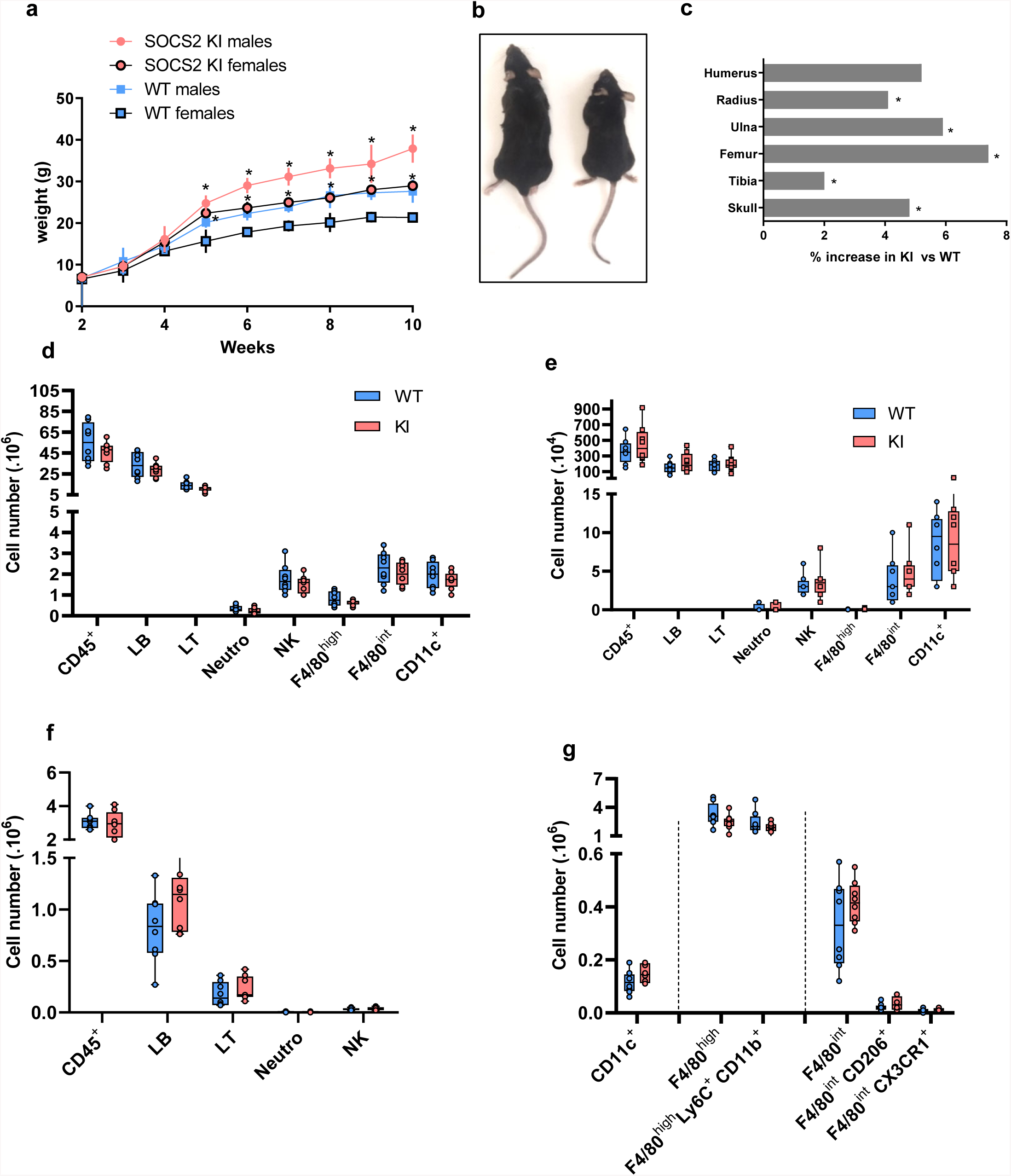
Characterization of SOCS2^R96C^ KI Mice. (**a**) Growth curves for male and female SOCS2^KI^ (circles) and WT (squares) mice. The body weight of the mice, weighed at weekly intervals, is shown. Each point represents the mean ± SD for 6-18 mice. (**b**) Increase in the size of a typical two-month-old SOCS2^KI^ male (left) relative to an age- and sex-matched wildtype animal (right). (**c**) Percentage increase in bone length in SOCS2^KI^ mice (right and left). Flow cytometry analysis of the immune landscape of single-cell suspensions from the (**d**) spleen, (**e**) lymph nodes, and (**f**) peritoneal cavity of two-month-old male SOCS2^KI^ or WT mice. N=8. (**g**) Flow cytometry analysis of macrophages/dendritic cell subsets from the peritoneal cavity of two-month-old male SOCS2^KI^ or WT mice, N=8. Statistical analysis was performed using the multiple t-test and significant p values are indicated. *P<0.05 vs. WT. SOCS, suppressor of cytokine signaling; WT, wildtype; KI, SOCS2^KI^ mice.

### SOCS2-R96C is not a direct downstream target of TLR ligation

Further investigation of the impact of the SOCS2-R96C mutation showed the number of bone marrow precursors to be significantly higher in SOCS2^KI^ mice, but the ratio of BM progenitors/body weight was significantly lower (Supplemental Figure 4). In mouse models, BMMs differentiated using M-CSF are a frequently used and convenient population to study macrophage function and signaling (Cunnick *et al*, 2006). We observed no differences in the phenotype between M-CSF derived macrophages from WT and SOCS2^KI^ mice. By flow cytometry analysis, both BMM cultures showed a homogeneous subpopulation of F4/80^+^ cells, with similar levels of CD11b and MHCII molecule expression (Supplemental Figure 5). Evidence has now accumulated indicating that SOCS1, SOCS3, and CIS are induced after TLR engagement in macrophages and DCs, and this contributes to avoid overshooting TLR stimulation (Dalpke *et al*, 2008). Similarly, it was shown (Hu *et al*, 2012) that various TLR ligands can induce SOCS2 gene expression in human DCs or thioglycolate-induced mouse peritoneal macrophages (Posselt *et al*, 2011). We thus investigated SOCS2 protein expression after TLR stimulation in WT and SOCS2^KI^ BMMs. Neither TLR-2 (FSL1, HKSA) nor TLR-4 (CRX) ligands induced SOCS2 expression in M-CSF-derived BMMs from WT or SOCS2^KI^ mice after 24 h of culture (Figure 2a) in contrast to previous reports using GM-CSF as a differentiating factor. Then, we investigated the direct effect of TLR ligands on pro-inflammatory cytokine production from BMMs from SOCS2^KI^ and WT mice after 24 h of culture. After 24 h of culture, there were no significant differences in IL-6 levels in cell culture supernatants between BMMs from WT and SOCS2^KI^ mice stimulated with FSL1, HKSA, or HKEB, regardless of the concentration of TLR ligand we used. Significantly higher IL-6 levels were observed only in the culture supernatants from SOCS2^KI^ BMMs after CRX stimulation at 10 nM relative to those from WT BMMs (Figure 2b). However, it is worth noting that FSL1stimulation induced higher production of GM-CSF by the BMMs from SOCS2-mutated than WT mice (Supplemental Figure 6).

**Figure 2.**
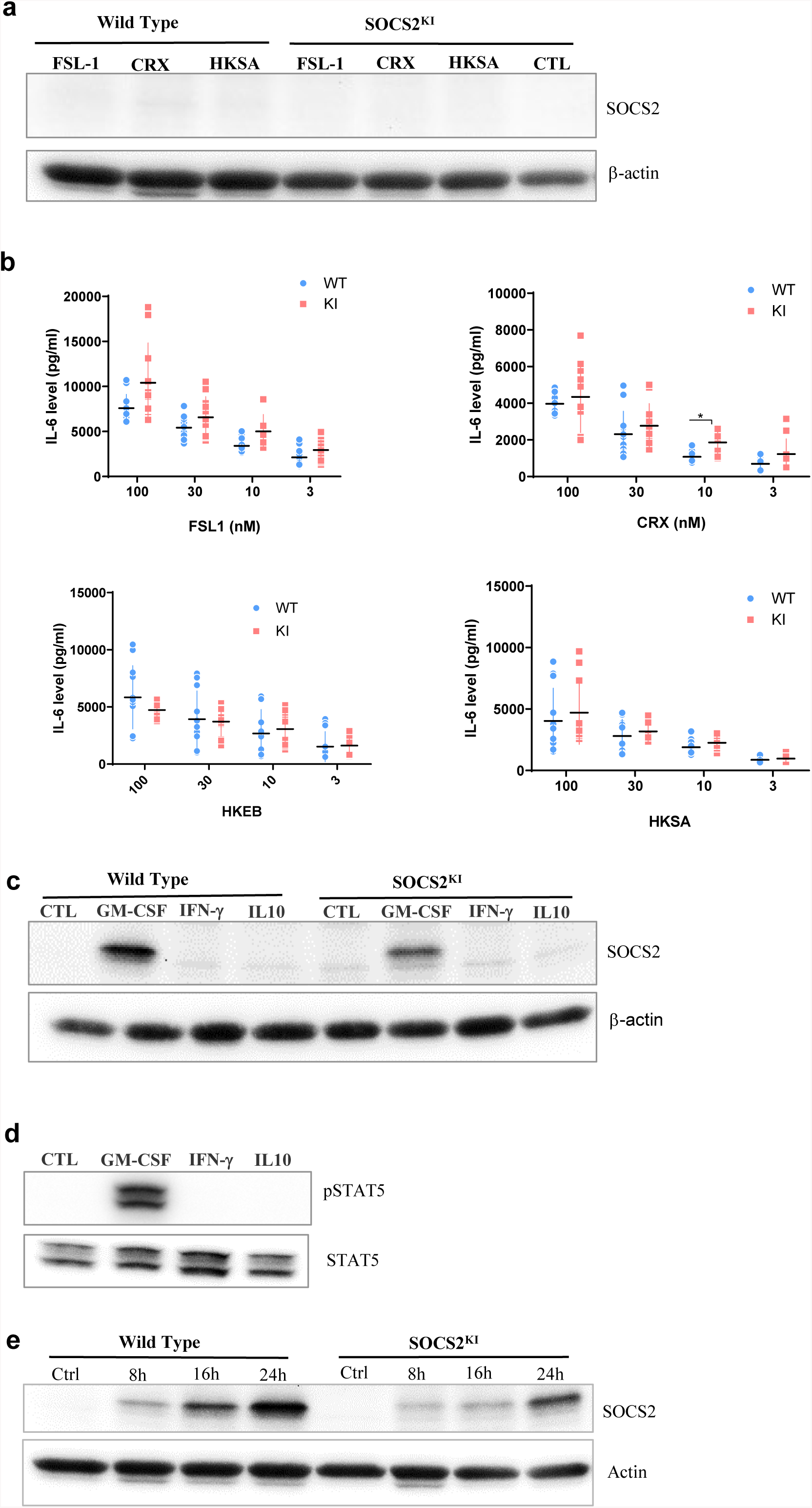
Response of BMMs from SOCS2^KI^ or WT mice to TLR ligands or cytokines. BMMs from SOCS2^KI^ or WT mice were stimulated 24 h with 3 to 100 nM of FSL1, CRX or HKSA or HKEB TLR ligands. (**a**) SOCS2 expression in cells was determined by western blotting. (**b**) IL-6 concentrations were measured in cell supernatants (Mean ± SD, N = 5). (**c**) BMMs from SOCS2^KI^ or WT mice were cultured with the cytokines GM-CSF, IFN-γ, or IL-10 at 30 ng/ml for 24 h and SOCS2 expression evaluated by western blotting. (**d**) BMMs from SOCS2^KI^ were cultured with the cytokines GM-CSF, IFN-γ, or IL-10 for 24 h and pSTAT5/total STAT5 expression were evaluated by western blotting. (**e**) The kinetics of SOCS2 expression were analyzed by western blotting after GM-CSF-culture of BMMs from SOCS2^KI^ or WT mice for 8, 16, and 24 h. Data are representative of two independent experiments. *P<0.05 vs. WT, by multiple group comparison of ANOVA. SOCS, suppressor of cytokine signaling; WT, wildtype; KI, SOCS2^KI^ mice.

SOCS2, a feedback inhibitor of JAK-STAT pathways, is well known to be upregulated in response to STAT-5-inducing cytokines, such as GM-CSF (Vitali *et al*, 2015; Rico-Bautista *et al*, 2006), whereas SOCS1 is induced by the cytokine IFN-γ via STAT-1 (Liu *et al*, 2020) and SOCS3 by IL-10 via STAT-3 (review in 27). We found SOCS2 protein expression in BMMs from WT mice to be induced by GM-CSF, but not IFN-γ or IL-10 stimulation. SOCS2 was also expressed by BMMs from SOCS2^KI^ mice at comparable levels after GM-CSF culture (Figure 2c). Moreover, as described in previous reports, we observed STAT-5 phosphorylation in WT BMMs after GM-CSF stimulation (Figure 2d). We next assessed SOCS2 expression from 8 to 24 h post-culture. In both BMMs cultures, SOCS2 expression continues to increase in WT but not in BMMs from SOCS2 KI mice. No difference was detected at 24 h, at which time similar levels between the two genetic backgrounds were measured (Figure 2e), indicating that SOCS2 regulates STAT-5 phosphorylation in a time-dependent manner during a short window of time after less than 24 h in BMM culture.

### Pro-inflammatory responses are enhanced in SOCS2^KI^ BMMs in relation to the cytokine context and the presence of GM-CSF

Macrophages secrete both pro-and anti-inflammatory cytokines following initial exposure to microbial products. As such, deletion of STAT-5 results in increased pro-inflammatory cytokine expression in mouse myeloid cells with PRR (pattern recognition receptor) expression (Brady *et al*, 2017). We explored the consequences of SOCS2-R96C mutation on pro-inflammatory cytokine secretion by culturing SOCS2^KI^ or WT BMMs with different concentrations of GM-CSF following FSL-1 or HKSA stimulation for 24 h. Cytokine concentrations are depicted as a heatmap according to the FSL-1 (from 3 to 100 nM) and GM-CSF concentrations (from 3 to 30 ng/ml) (Figure 3). High concentrations of FSL-1 (100 or 30 nM) in culture, even with a low concentration of GM-CSF (3 ng/ml), led to greater secretion of IL-6 by SOCS2^KI^ mice than WT mice (Figure 3a). Indeed, stimulation with FSL1 (at 100 and 30 nM) resulted in significantly higher production of IL-6 by BMMs from SOCS2^KI^ mice (4-fold increase) (Figure 3). Interestingly, TNF-α secretion was not affected by the loss of SOCS2 function (Figure 3b).

**Figure 3.**
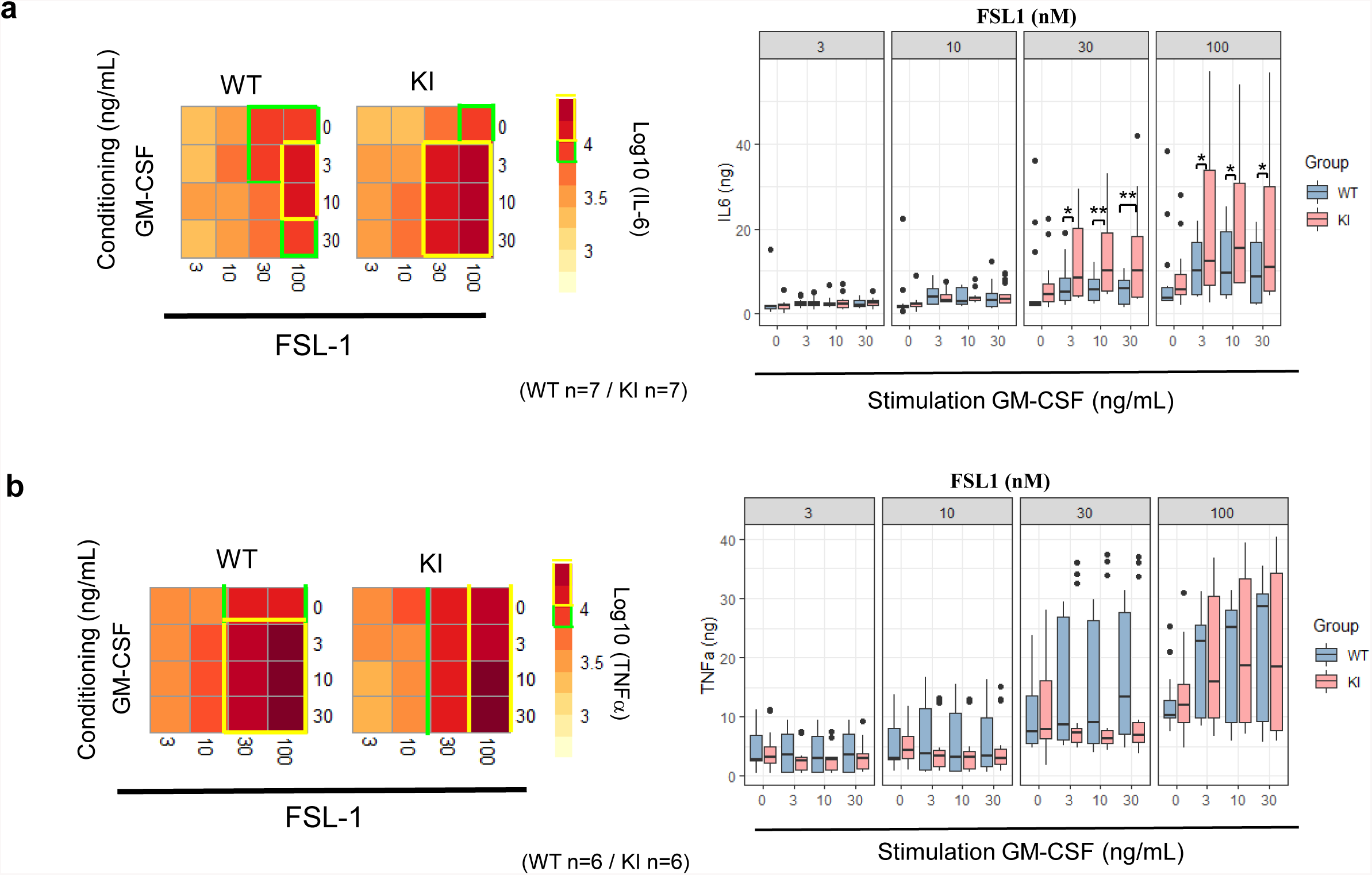
IL-6 and TNF-α response to FSL1 in GM-CSF-conditioning medium. Heatmap representation of IL-6 (**a**) and TNF-α (**b**) concentrations in supernatants of BMMs from SOCS2^KI^ or WT mice (after Z-score transformation) after culture with increasing concentrations of FSL1 (from 3 to 100 nM) and GM-CSF (from 3 to 30 ng/ml). IL-6 (**a**) and TNF-α (**b**) concentrations (pg/mL) in BMM supernatants after culture with increasing concentrations of FLS1 and GM-CSF. Results represent the mean ± SD of 6-7 independent donors. *P<0.05, **P<0.01, ***P<0.001, ****P<0.0001 vs. WT, by multiple group comparison of ANOVA.

### STAT-5 phosphorylation is sustained in SOCS2^KI^ BMMs after TLR stimulation

It has been shown that SOCS negatively regulates cytokine signaling through various mechanisms that target the JAK/STAT pathway (Croker *et al*, 2008). We thus studied the kinetics of STAT-5 phosphorylation by inducing SOCS2 expression by a pulse of GM-CSF for 24 h followed by a 3-h chase. Then, pSTAT-5 levels were monitored from 10 to 120 min after re-introduction of GM-CSF to the BMM culture (Figure 4a). We observed rapid phosphorylation of STAT-5 after 10 min of stimulation in both genetic backgrounds. However, STAT-5 phosphorylation was sustained in BMMs from SOCS2^KI^ mice for up to 30 min post-stimulation, whereas it decreased in WT cells (Figure 4b). Relative to WT mice, STAT-5 dephosphorylation was significantly delayed in SOCS2^KI^ mice, consistent with our working hypothesis of a difference in regulation of the JAK/STAT pathway.

**Figure 4.**
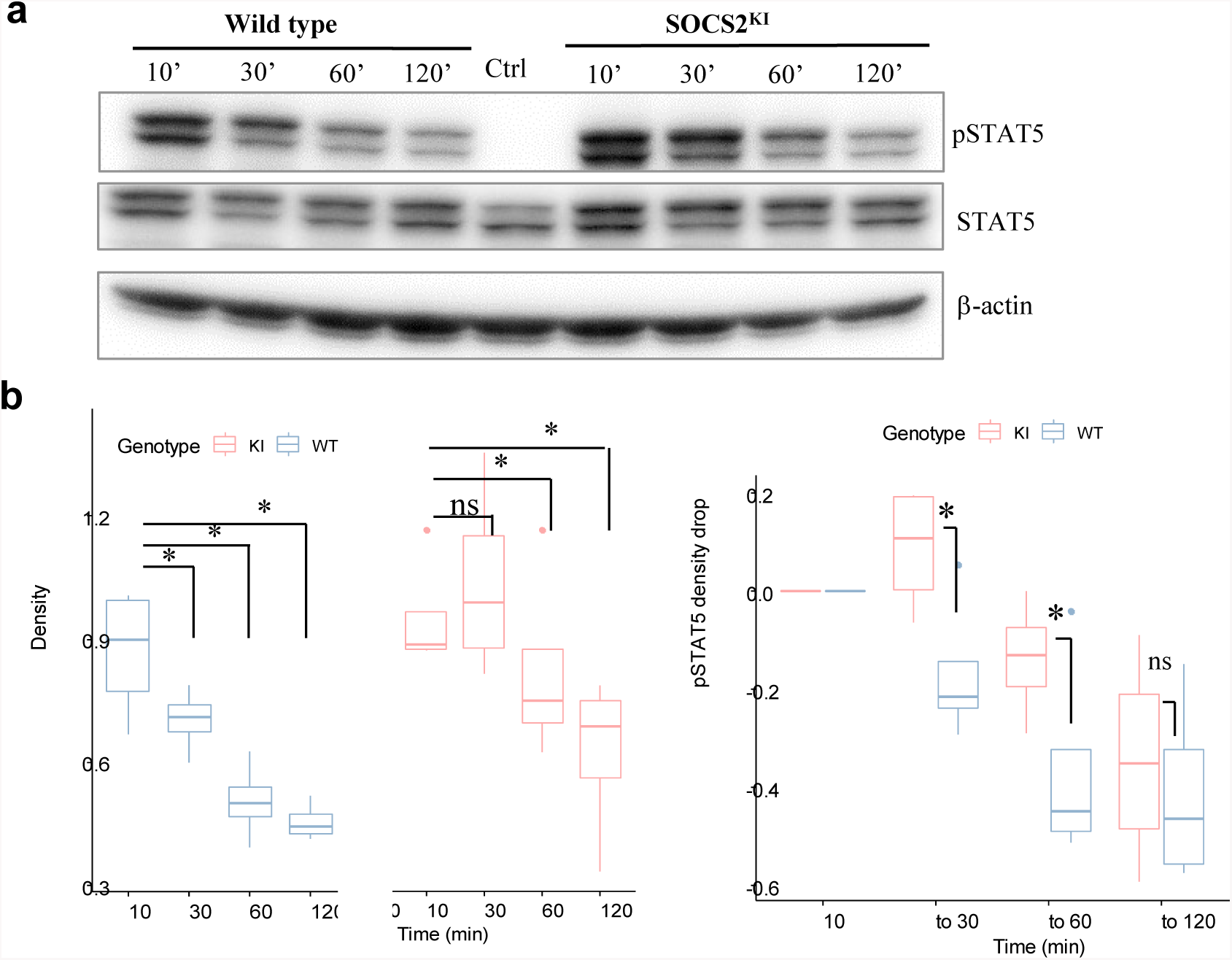
Kinetics of pSTAT5 expression in GM-CSF-cultured BMMs from SOCS2^KI^ or WT mice. BMMs were cultured in the presence of GM-CSF (10 ng/ml) for 24 h followed by a chase of 3 h in culture medium without cytokines. (**a**) Expression of pSTAT5 was analyzed after the re-addition of GM-CSF at 10 ng/ml for 10 min to 2 h of stimulation by western blotting. (**b**) Density representation of pSTAT5 blots as a function of duration of stimulation with GM-CSF for WT and SOCS2^KI^ BMMs. (**c**) Drop in pSTAT5 density. **P<0.01, Kruskal Wallis test.

### SOCS2 regulates SOCS1 but not SOCS3 cytokine responses in a dose-dependent manner

Previous studies indicated that SOCS2 may regulate other SOCS proteins, in particular SOCS1 and SOCS3 (Tannahill *et al*, 2005), in a dose-dependent manner (Dif *et al*, 2001). Through this mechanism, SOCS2 can indirectly modulate cytokine-induced STAT activation by removing the negative regulation mediated by the SOCS proteins that are targeted. Indeed, it is generally considered that IFN-γ mainly induces SOCS1 via STAT-1, whereas IL-10 induces SOCS3 via STAT-3 (Sobah *et al*, 2021).

We investigated the role of SOCS2 on SOCS1- and SOCS3-regulated cytokine responses by priming BMMs with IFN-γ and IL-10 before stimulation with the FSL-1 ligand. The cytokine response and phagocytic ability of cytokine-sensitized macrophages were quantified. First, IL-6 levels were significantly higher in culture supernatants from SOCS2^KI^ BMMs than those from WT BMMs after 30 nM of FSL-1 stimulation in the presence of various concentrations of IFN-γ and IL-10 (from 3 to 30 ng/ml) (Figure 5a). BMMs from the KI mice also produced higher levels of TNF-α but only with IL-10 conditioning medium, even at the low FSL-1 concentration (3 nM) (Figure 5b). By contrast, phagocytosis was not significantly different between the genetic backgrounds or in the presence of the cytokines GM-CSF, IL-10, or IFN-γ used to sensitize the BMMs (Supplemental Figure 7).

**Figure 5.**
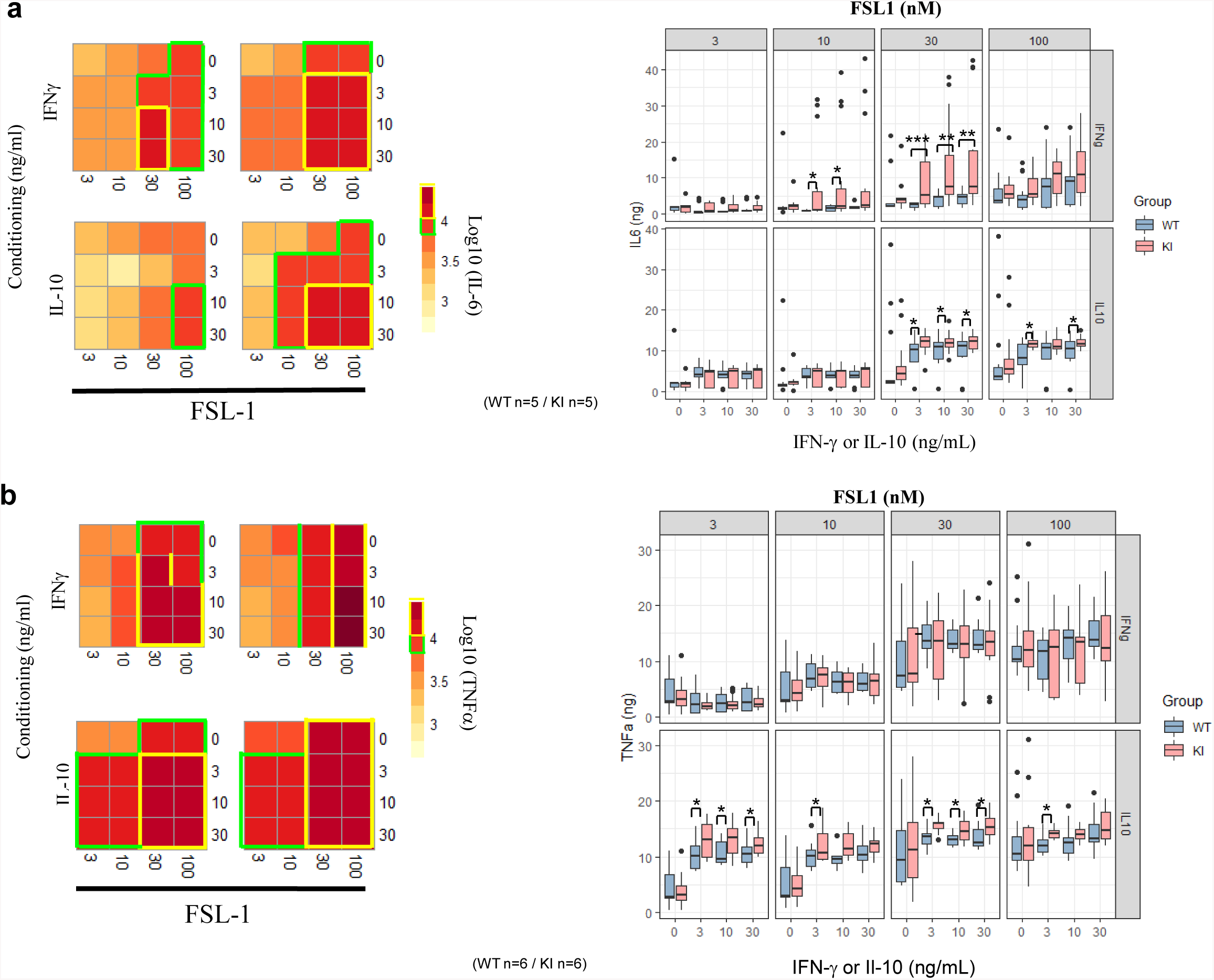
IL-6 and TNF-α response of BMMs to FSL1 in IFN-γ and IL-10 conditioning medium. Heatmap representation of IL-6 (**a**) or TNF-α (**b**) concentrations in supernatants of BMMs from SOCS2^KI^ or WT mice (Z-score transformation) after culture with increasing concentrations of FSL1 (3 to 100 nM) and IFN-γ or IL-10 (3 to 30 ng/ml). IL-6 (**a**) and TNF-α (**b**) concentrations (pg/mL) in BMM supernatants after culture with FLS1 (3 to 100 nM) and IFN-γ or IL-10 (3 to 30 ng/ml). Results represent the mean ± SD of 5-6 independent donors. *P<0.05, **P<0.01, ***P<0.001, ****P<0.0001 vs. WT, by multiple group comparison of ANOVA.

As IL-6 was differentially secreted by BMMs from mice of the two backgrounds, we sought differences in signaling focusing on the gp130-containing IL-6 receptor family. SOCS3 has already been shown to negatively regulate STAT-3 activation in response to several cytokines, such as those of the gp130-containing IL-6 receptor family (Carow & Rottenberg, 2014). We investigated the indirect impact of the SOCS2-R96C mutation on STAT-3 activation after IL-6 stimulation from 10 to 60 min. There were no differences in pSTAT-3 levels between SOCS2^KI^ and WT BMMs, regardless of the time of stimulation (Supplemental Figure 8).

### The loss of SOCS2 function changes the immune response to *S. aureus* and worsens the outcome of the infection

We further investigated the impact of the R96C mutation on the outcome of infection with *S. aureus* using a peritonitis model (Kim *et al*, 2014) (Figure 6a). After 16 h of infection, leukocyte numbers in peritoneal exudates from SOCS2^KI^ mice were significantly higher than those from WT mice. This increase was associated with a higher number of neutrophils (Ly6G^+^ cells) and macrophage subsets (F4/80^int/+^ cells). PLSDA analysis showed a clear dichotomy between the two genotypes using PerCs recruitment data (Figure 6b). *Ex vivo* imaging showed higher neutrophil recruitment in SOCS2^KI^ than WT mice. Interestingly, injection of heat-killed *S. aureus* (HKSA) failed to recapitulate the observed inflammatory response pattern (Supplemental Figure 9). After 48 h of *S. aureus* infection, the dynamics of inflammatory cell recruitment were highly similar between the WT and mutant mice (Figure 6c). As R96C mutation was associated with an increase in cell number in the neutrophils and macrophages subsets at early stages of infection, we studied bacterial fitness after infection. In accordance with the increase in the number of neutrophils, there was a significantly lower number of total live bacteria 16 h post infection (Figure 7a), associated with an increase in efferocytosis (increase in the number of F4/80^+^Ly6G^+^ cells) (Figure 7b). Interestingly, the peritoneal exudates of SOCS2^KI^ mice contained much more IFNγ and CXCL10 than those of WT mice (Figure 7c). By contrast, the total number of live bacteria was significantly higher in SOCS2^KI^ mice than WT mice 48 h post infection, indicating that, despite early recruitment of inflammatory cells, the infection was not contained in the SOCS2^KI^ mice and continued to progress.

**Figure 6.**
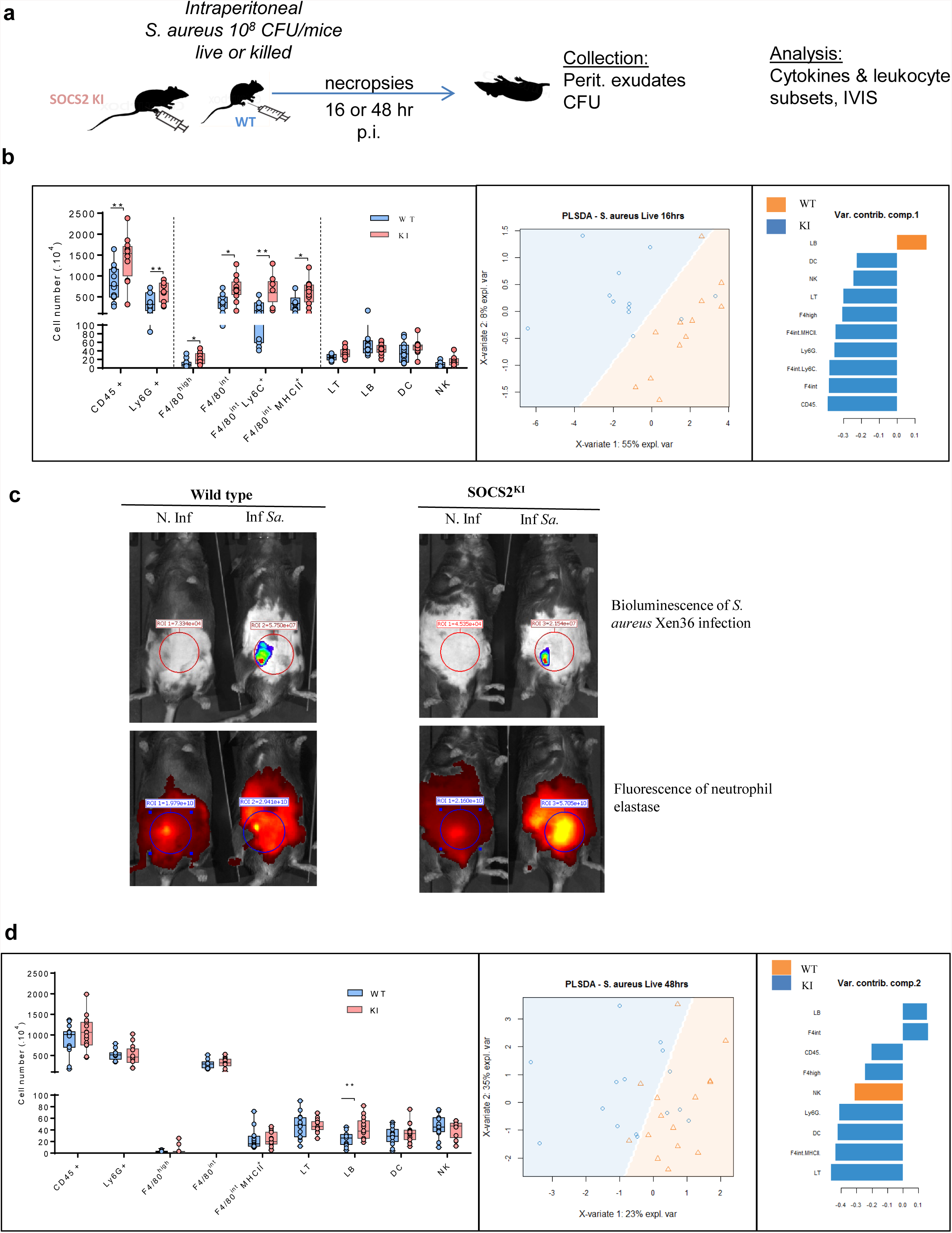
Immune cell analysis after 16 or 48 h of *S. aureus* peritonitis in SOCS2^KI^ or WT mice. (**a**) Experimental design: male SOCS2^KI^ or WT mice were peritoneally challenged with the *S. aureus* Xen36 strain at 10^8^ CFU/mice (N=12). (**b**) Immune cell-composition in peritoneal cavity analyzed by flow cytometry 16 h after infection. PSLDA showing hierarchical clustering of individual mice as a function of immune cell-composition and the respective contribution of the quantitative variables to dimensions 1 or 2. (N=12) (**c**) *Ex-vivo* imaging of luminescence (*S. aureus*) and fluorescence intensity of elastase (neutrophil) 16 h post infection. Data are representative of two independent experiments. (**d**) Immune cell-composition 48 h post infection in the peritoneal cavity analyzed by flow cytometry and PSLDA clustering (N=12). *P<0.05, **P<0.01, by multiple group comparison of ANOVA.

**Figure 7.**
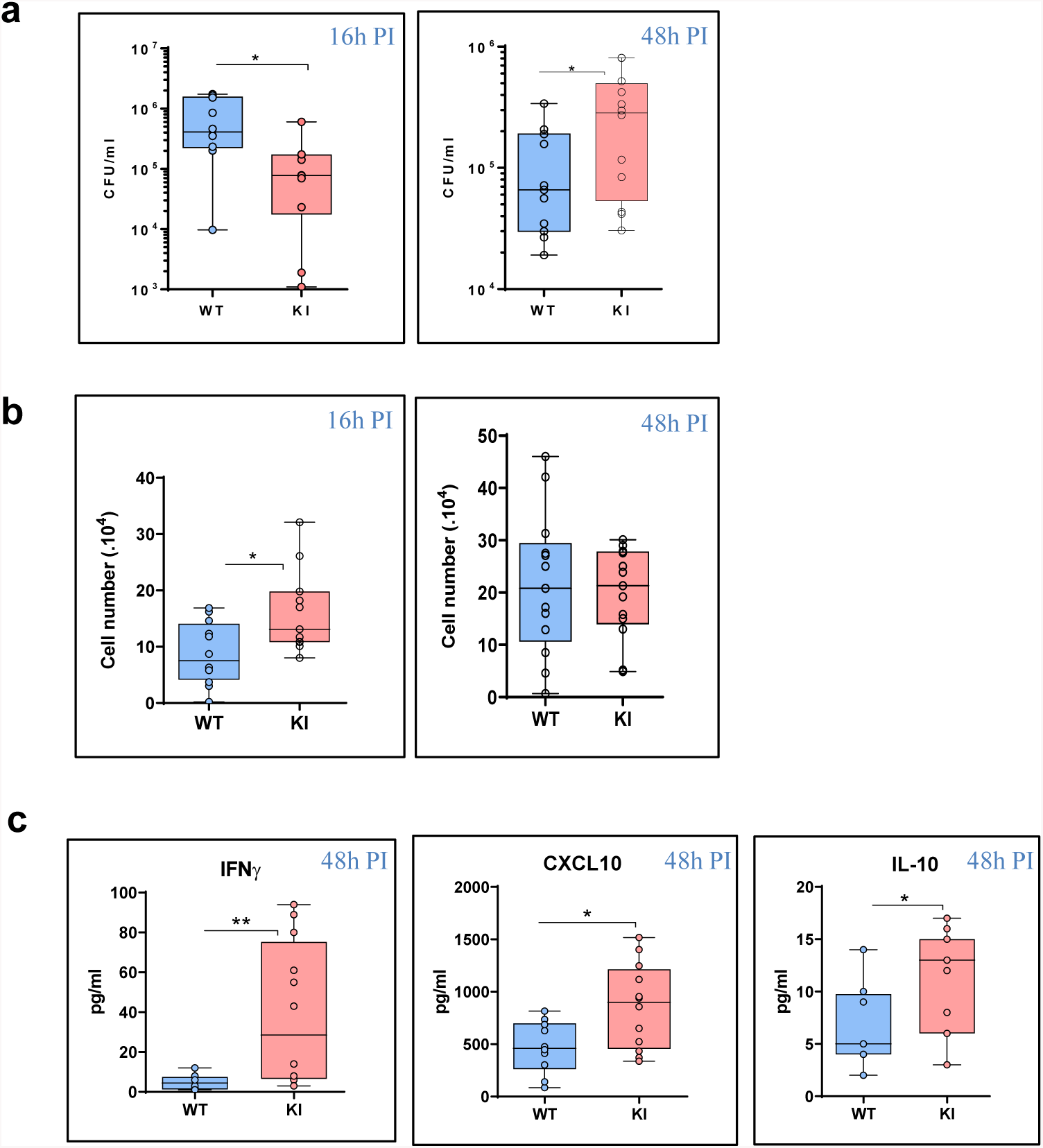
Analysis of bacterial fitness and the immune response after 16 or 48 h of *S. aureus* peritonitis in SOCS2^KI^ or WT mice. (**a**) Total live bacteria in the peritoneal lavage 16 and 48h post infection. (**b**) Efferocytosis in the peritoneal cavity 16 and 48h post infection (corresponding to total F4/80^+^Ly6G^+^ cells/live CD45^+^ cells). (**c**) Cytokine concentrations (pg/mL) in the exudates determined by multiplexed ELISA. Results are represented by box and whisker plots showing the Min to Max (N=12). Statistical analysis was performed using the Mann-Whitney test and significant p values are indicated. *P<0.05 vs. WT.

## DISCUSSION

Here, we provide evidence that the loss-of-function R96C mutation of the SOCS2 protein considerably alters orchestration of the inflammatory response both *in vitro* and *in vivo*. Macrophages from the R96C mutant genetic background showed heightened responsiveness to TLR ligands in terms of STAT activation and cytokine production, which characterize activated macrophages. Furthermore, *in vivo* studies using a *S. aureus* peritoneal infection model show that SOCS2-R96C is associated with immune dysregulation, with elevated recruitment of macrophage subsets and neutrophils at early stages and increased expression of pro-inflammatory cytokines and chemokines.

SOCS2 has two major functional domains: a central SH2 domain and a SOCS box. The SOCS2-SH2 domain directly interacts with JAK2, whereas the SOCS box mediates proteasomal degradation. Recently, Li et al. (2022) (Li *et al*, 2022) showed that the R96C mutation in the SH2 domain of SOCS2 in mice does not compromise the domain integrity of the SOCS box nor alter the ability of SOCS2 to recruit the SOCS box-associated-E3 ubiquitin ligase complex. Here, we show that SOCS2^R96C^ KI mice display a similar increase in growth as previously reported for *Socs2^-/-^* mice, with an identical increase in body and organ weight and bone length (Metcalf *et al*, 2000; Greenhalgh *et al*, 2002). Overall, these observations confirm that the SOCS2^R96C^ KI mouse model provides a new tool to evaluate the functions of SOCS2 using a protein with a deficient SH2-domain. Indeed, this model, unlike the *Socs2*^-/-^ model, makes it possible to study the interactions between proteins, in particular the contribution of SH2:pTyr binding to the function of SOCS2 in immune responses, while the expression of the SOCS2 protein is maintained, lowering the risk of perturbing interactive networks involving SOCS proteins.

An increasing number of studies suggest that SOCS proteins participate in pattern recognition receptor (PRR) signaling. SOCS2 is a cytokine-inducible inhibitor of the JAK-STAT signaling pathway, with several studies reporting that SOCS2 is directly induced by TLR ligation in DCs and macrophages (Machado *et al*, 2006; Hu *et al*, 2012). Hu et al., (2012) (Hu *et al*, 2012) showed that various TLR ligands induce *Socs2* gene expression in human DCs. More precisely, TLR-4 signaling in monocyte-derived DCs induces the production of type I interferon, which in turn activates SOCS2 via STAT-3 and STAT-5. Moreover, Posselt et al. (2012) (Posselt *et al*, 2011) demonstrated that LPS stimulation of thioglycolate-induced mouse peritoneal macrophages results in significant upregulation of the mRNA levels of *Socs2*, which regulates IL-1beta and IL-10. The same group showed that *Socs2* mRNA is induced in monocyte-derived DCs upon TLR-8 and NOD signaling, thus controlling the release of pro-inflammatory mediators from DCs (Schwarz *et al*, 2013).

Our results are not in accordance with these findings. Indeed, BMM stimulation for 24 h with multiple TLR agonists failed to induce SOCS2 expression at the protein level. This observation suggests the absence of any direct relationship between TLR-triggered pathways and SOCS2 expression. Rather, we believe that certain cytokines produced as a result of TLR stimulation may bind to their own receptor in an autocrine fashion and possibly induce SOCS2 expression. For example, we found that SOCS2^KI^ BMMs produce higher quantities of GM-CSF after FLS-1 stimulation.

Macrophages are key immune cells that play a crucial role in regulating and maintaining tissue homeostasis (Lavin *et al*, 2015), as well as in immune defense against invading pathogens. Macrophages generated from bone-marrow progenitors or monocytes in the presence of GM-CSF are thought to be typically M1-like, producing pro-inflammatory cytokines upon stimulation with TLR ligands (Murray *et al*, 2014). The binding of GM-CSF to its receptor induces STAT-5 phosphorylation, which is tightly regulated by SOCS2 (Review in 40,41). Consistent with this observation, we found that only GM-CSF, in contrast to the cytokines IFN-γ and IL-10, induced STAT-5 phosphorylation and SOCS2 expression in BMMs from both WT and SOCS2^KI^ mice. Moreover, we found that the R96C mutation resulted in sustained STAT-5 phosphorylation after GM-CSF stimulation. Moreover, Zhan et al. (2019) (Zhan *et al*, 2019) recently reported that the amount of GM-CSF is a key factor in determining its biological activity. We investigated whether SOCS2 could have a direct activity on TLR-induced cytokine secretion by conditioning BMMs from both genotypes with increasing amounts of GM-CSF and further stimulating them with the TLR-2 agonist FSL-1.

Our results showed higher pro-inflammatory IL-6 production by SOCS2^KI^ BMMs that was dependent on the GM-CSF and FSL-1 ligand concentrations. Interestingly, the specificity of the response was highly specific as a function of the type of pro-inflammatory cytokine (IL-6 vs. TNF-α). It is well known that SOCS1 production is induced by the cytokine IFN-γ via STAT1 (Liu *et al*, 2020) and that of SOCS3 by IL-10 via STAT3 (review in 27). Acceleration of SOCS3 degradation has been observed in cell lines overexpressing SOCS2 (Tannahill *et al*, 2005; Piessevaux *et al*, 2006). By contrast, Kiu et al. (2009) (Kiu *et al*, 2009) showed that SOCS2 is not a physiological regulator of SOCS3 expression or its action in primary hematopoietic cells using *Socs2*^-/-^ mice. Previous work showed that the loss of STAT-5 in hepatocytes results in reduced expression of its target genes *Socs2* and *Socs3*, which leads to exacerbated STAT3 signaling through gp130-based receptors (Cui *et al*, 2007). Our results show that increasing the IL-10 concentration provokes significantly stronger pro-inflammatory cytokine responses by SOCS2^KI^ BMMs after TLR-2 engagement than by WT BMMs. We obtained comparable results in IFN-γ conditioning medium, although the cytokine response appeared to be dependent on the TLR-ligand used. Previous studies have also shown that *Socs2* expression is upregulated by IFN-γ in DCs in human melanoma (Nirschl *et al*, 2017). Such data suggest that SOCS2 could negatively regulate SOCS3 protein levels and thus be a positive modulator of SOCS3-inhibited cytokine signaling cascades.

Here, we elucidated the role of R96C-SOSC2 in an experimental model of *S. aureus* peritonitis. SOCS2^KI^ mice showed early intense myeloid cell recruitment, with elevated numbers of neutrophils and inflammatory macrophages in the peritoneal cavity associated with a decrease in bacterial fitness. Moreover, *S. aureus* peritonitis in SOCS2^KI^ mice resulted in higher IFN-γ, CXCL10, and IL-10 levels than in WT mice. Consistent with this observation, *Socs2*^-/-^ mice show uncontrolled Th1 cell-mediated responses to *Toxoplasma gondii*; leading to death, suggesting an increased proinflammatory response. SOCS2^KI^ mice showed an increase in macrophage efferocytosis at the early stages of infection, without a modification of phagocytosis. On the contrary, in an acute arthritis model, *Socs2*^-/-^ macrophages exhibited reduced efferocytosis, but only for large peritoneal macrophages (F4/80^high^) (Cramer *et al*, 2022). These authors concluded that SOCS2-deficient mice exhibit less apoptosis and efferocytosis than WT mice at the late adjuvant-induced arthritis phase (Cramer *et al*, 2022). At the late stage of *S. aureus* infection, SOCS2^KI^ mice showed a higher bacterial load, associated with a higher number of B lymphocytes. Our results strongly suggest that SOCS2 is required for the late resolution of the inflammatory response to bacterial infection.

Our results show that the SOCS2 protein plays an important role in the immune response by controlling inflammatory cytokine production and reducing cell infiltration at the early stage of infection. These results suggest that SOCS2 highly contributes to the recruitment process of immune cells and the regulation of the production of various pro-inflammatory cytokines during a bacterial infection.

## MATERIALS & METHODS

### Mouse strains

The study was carried out in compliance with the ARRIVE guidelines (http://nc3rs.org.uk/arrive-guidelines). The R96C point mutation was introduced into the C57Bl/6 mouse genome by homologous recombination using CRISPR/Cas9 technology. One single nucleotide was replaced by homologous recombination in pronuclear-stage zygotes, as previously described (Pelletier *et al*, 2015). The C57Bl/6 background was chosen to facilitate the production of further recombinant mice (gene reporter or knockout strains) of interest to elucidate the mechanisms altered by the point mutation. Eight- to 10-week-old female or male WT C57BL/6 mice or SOCS2^KI^ mice were bred and housed in a specific pathogen-free facility (INSERM US 006 – CREFRE). Body growth was determined for female and male WT and SOCS2^KI^ mice. Experiments were performed in an accredited research animal facility of the UMR IHAP, ENVT, Toulouse, France. Mice were handled and cared for according to the ethical guidelines of our institution (APAFIS#22936-2019112515186332) following the Guide for the Care and Use of Laboratory Animals (National Research Council, 1996) and in compliance with European directive 2010/63/UE under the supervision of authorized investigators. Mice were euthanized by cervical dislocation and all efforts were made to minimize pain and distress of the animals.

### Computed tomography imaging of living mice

Anesthetized (Isoflurane, Virbac) 10-week-old mice (n=6) were scanned using a small animal computed tomography (CT) system (NanoScan PET/CT Mediso). The nanoScan CT has a rotating gantry and the X-ray source and detector rotate around the object. The scanning protocol was 35 kVp, 800 µA, 450 ms integration time, 720 projections per 360°, scan duration 5’33”. The reconstructed voxel size was 125 x 125 x 125 µm. From the acquired data, the length of the humerus, radius, ulna, femur, and tibia and the length and width of the skull were bilaterally determined. To ensure reproducibility of the measurements, precise anatomical landmarks were defined beforehand.

### Preparation of *S. aureus* and challenging of the mice

*Staphylococcus aureus subsp. aureus (ATCC® 49525™)* XEN36 (bioluminescent bacterial strain) or the HG001 strain, a GFP-expressing mutant (Herbert *et al*, 2010), were grown overnight in TSB at 37°C with orbital shaking (200 rpm). The culture was further diluted 1:100 in TSB and grown to mid-log phase (O.D. 600 nm _1). Bacterial cells were pelleted (2800 x g, 10 min, 4°C), washed twice in PBS, and resuspended in physiological serum. The concentration of the bacteria was estimated by measuring the absorbance at 600 nm (OD600=0.8 for 2×10^8^ CFU/mL) and confirmed by serial dilution in PBS Tween 20 (0.05%) and plating on LB-Agar for determination of the number of colony-forming units (CFU). CFUs were determined after 24 h of incubation at 37°C. Bacteria were freshly prepared before each experiment and adjusted to the desired concentration. The XEN36 strain was killed by heating for 1 h at 65°C. *S. aureus* inactivation was confirmed by LB culture. Mice were infected intraperitoneally (i.p.). with *S. aureus*. at 10^8^ CFU/mouse (100 ul/mouse).

### Peritoneal cell (PerC) collection and spleen and lymph node cell isolation

The peritoneal cavity was washed by injecting 3 mL PBS. After recovery, peritoneal exudates were centrifuged (300 x g, 5 min) and the supernatants stored at −80°C for further analysis. PerCs were harvested for cell phenotyping by flow cytometry. The spleens and lymph nodes (LNs) were removed and the cells isolated through 70 and 40 μm cell strainers, respectively, to obtain single-cell suspensions in PBS.

### Flow cytometry analysis

The number of cells obtained from the peritoneal exudates, spleens, and LNs was determined using a flow cytometry absolute counting system (MACSQuant Analyzer, Miltenyi Biotec, Germany). Cells (1-2 x 10^6^) were incubated in HBSS, 0.5% BSA, 10 mM Hepes containing mouse FcR Blocking Reagent (Miltenyi Biotec, Germany) following the manufacturer’s instructions. Cell viability was assessed using Viobility 488/520 Fixable Dye (Miltenyi Biotec, Germany). Antibodies were incubated at 4°C for 30 min in the dark. The antibodies used are listed in Supplemental Table 1 (Supplementary Information). Flow cytometry was used to measure efferocytosis by quantifying the number of F4/80^int^ Ly6G^+^ macrophages in the peritoneal cavity from WT or SOCS2^KI^ mice after *S. aureus* infection. Acquisition was performed using a MACSQuant (Miltenyi Biotec, Germany) flow cytometer with MACS Quantify software. Flow cytometry data were analyzed using FlowJo (Tree Star, USA) software.

### Preparation of mouse bone marrow-derived macrophages (BMMs)

Femurs and tibias from WT or SOCS2^KI^ mice were cut at both ends and the bone marrow flushed out with PBS with a syringe mounted with a 26-gauge needle. The cells were collected and cultured in X-Vivo (BE02-060F, Ozyme, France) supplemented with 50 ng/mL M-CSF (Peprotech, France) at a density of 2 x10^5^ cells/mL. Cells were incubated at 37°C in 5% CO_2_ and fresh medium added on days 3 and 5 of culture. On day 7, the BMMs were harvested, counted, and primed for 24 h with either GM-CSF (Biolegend, France), IFN-γ (Peprotech, France), or IL-10 (Peprotech, France) at 3, 10, or 30 ng/ml. The TLR ligands FLS1 (Invivogen, France) and CRX (Invivogen, France) at various concentrations (3, 10, 30, or 100 nM) or heat-killed *S. aureus* (HKSA) (Invivogen, France) or heat-killed *E. coli* (HKEB) (Invivogen, France) at various MOIs (3, 10, 30, or 100) were added to the cell cultures and the cultures incubated for an additional 24 h. We used the mycoplasma lipoprotein FSL-1, recognized by the TLR2/TLR6 heterodimer, HKSA, recognized mainly by TLR2, CRX-527, a highly specific TLR4 agonist, and an LPS-like molecule and HKEB recognized by TL2 and TLR4. Cell supernatants were then harvested for cytokine detection. BMMs were pulsed 24 h with GM-CSF at 10 ng/ml to induce SOCS2 expression followed by a 3 h chase. Then, GM-CSF (10 ng/ml) was added de novo to cells followed by incubation for 10, 30, 60, or 120 min. Cells were harvested to analyze SOCS2 and pSTAT5/total STAT5 expression by western blotting.

### Measurement of cytokine/chemokine production

Cytokines in peritoneal exudates were quantified using a customized multiplex assay kit containing IL-1β, IL1-α, IL-10, IL-6, IFN-γ, IL-12, IL-17, CXCL1, CXCL10, CCL2, CCL3, CCL4, and TNF-α (Milliplex-MAP, Merck Millipore, France) and a xMAP instrument (MAGPIX, Luminex). Individual cytokine detection kits were also used to quantify mouse IL-6 and TNF-α (Bio-techne, France) and GM-CSF (Biolegend, San Diego, USA).

### Western-blot analysis for SOCS2 protein or related nuclear transcriptional factors

After stimulation, BMMs were washed once with cold PBS. Total protein was extracted with radio immune precipitation assay (RIPA) buffer containing phenylmethylsulphonyl fluoride (PMSF) (Roche, Switzerland) and a protease inhibitor cocktail (Roche) before storage at −80°C. Protein concentrations were determined using the BCA method. Cell lysates (30 µg per lane) were subjected to SDS polyacrylamide gel electrophoresis and then transferred to polyvinylidene difluoride membranes for western-blot analysis. After blocking with 5% fat-free milk dissolved in TBS-T (Tris buffered solution with Tween 20) for 1 h at room temperature, membranes were incubated overnight with antibodies raised against total STAT3, pSTAT3, pSTAT5, total STAT5, and SOCS2 (Cell Signaling Technology, Beverly, MA) according to the manufacturer’s instructions. Binding of these primary antibodies was visualized using goat anti-rabbit/anti-mouse immunoglobulin coupled to horseradish peroxidase (Jackson ImmunoResearch). The measurement of β-actin served as a loading control.

### Phagocytosis assay

To assess phagocytosis, BMMs were first primed with either GM-CSF (Biolegend, France), IFN-γ (Peprotech, France), or IL-10 (Peprotech, France) for 24 h and then infected at a MOI of bacteria to macrophages of 10:1 with *S. aureus* HG001-GFP for 1 h at 37°C or 4°C as a control. Each well was washed three times and extra-cellular fluorescence quenched by adding 0.2 uM syringe-filtrated trypan blue (Sigma). Cell viability was determined using 7-AAD dye (Biolegend). The amount of live (GFP^+^ 7AAD^-^) and dead (GFP^+^ 7AAD^+^) engulfed bacteria was measured by flow cytometry (MACSQuant, Miltenyi Biotech, Germany).

### *Ex vivo* bioimaging

Mice were infected i.p. with bioluminescent *S. aureus* XEN36 at 10^8^ CFU/mouse (100 ul/mouse). A neutrophil elastase solution (680 FAST, Perkin Elmer) was injected intravenously (4 nmol/100 ul/mouse) 3 h later. The animals were imaged by bioluminescence and fluorescence 24 h after *S. aureus* infection.

### Quantification and statistical analysis

Prism v. 8.0.1 (GraphPad software, San Diego, CA, USA) was used for all analyses. Significant differences were analyzed using the t-test or multiple t-test of ANOVA. P<0.05 was considered statistically significant. Statistical computing was carried out and some of the graphics generated using R software. Graphics were designed using the ggplot2 R package. Heatmaps were computed after transformation of the cytokine concentrations into Z-scores and generated using the Pheatmap R package (Kolde Raivo). PLSDA was carried out using the MixOmics package (Rohart *et al*, 2017).

## ACKNOWLEDGEMENTS

This work was supported by a grant from the Agence Nationale de la Recherche (ANR REIDSOCS; ANR-16-CE20-0010).

## AUTHOR CONTRIBUTIONS

LG, BG, GT, and CH designed the experiments, analyzed the data, and wrote the manuscript. LG, CH, CT, and BG performed the research and prepared the figures. All authors contributed to the article and approved the submitted version.

## DECLARATION OF INTERESTS

All authors declare no competing interests

## SUPPLEMENTAL FIGURES

**Suppl Figure 1.**
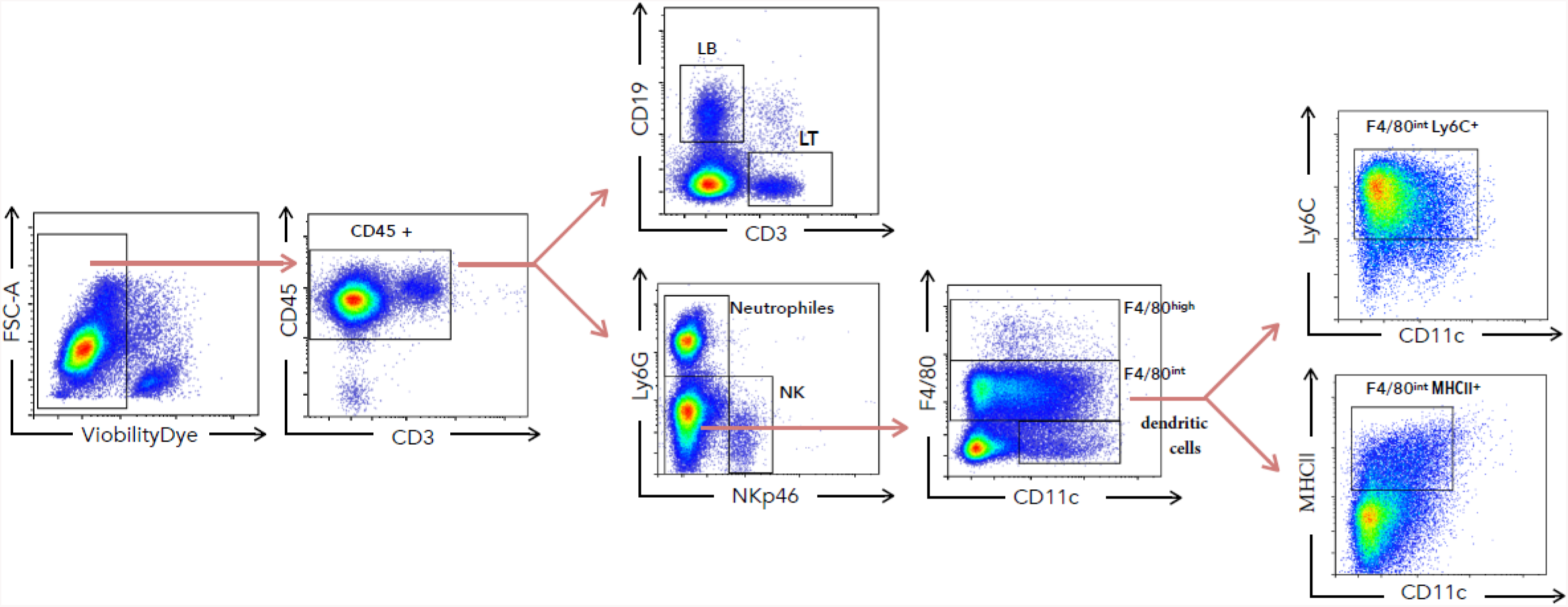
Flow cytometry gating strategy. Flow cytometry gating strategy for the labeling and analysis of immune-cell sub-populations in peritoneal exudates. Leukocytes (identified as CD45^+^), were divided among B cells (CD19^+^), T cells (CD3^+^), neutrophils (Ly6G^+^), NK cells (NKp46^+^), dendritic cells (CD11c^+^), and two sub-populations of macrophages: resident macrophages (F4/80^high^) and inflammatory macrophages (F4/80^int^). This inflammatory macrophages were further separated into CD11c^low^/MHCII^+^ cells and CD11c^low^ Ly6C^+^ cells.

**Suppl Figure 2.**
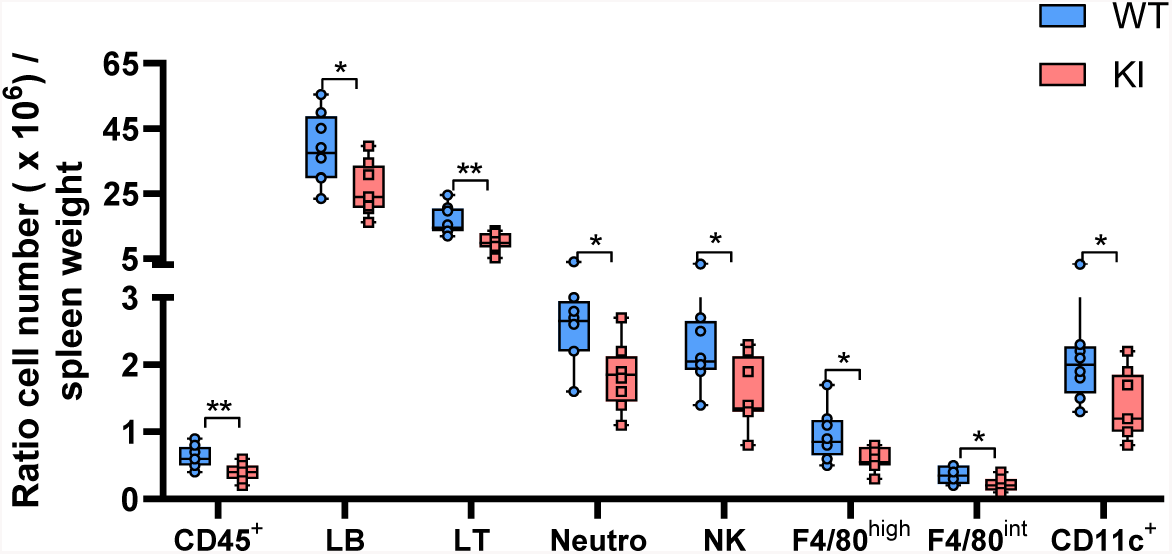
Ratio of cell number/weight of spleens from SOCS2^KI^ or WT mice. The ratio of the cell number/weight of spleens from two-month old male SOCS2^KI^ or WT mice (N=8). Statistical analysis was performed using the multiple t-test of ANOVA. *P<0.05 vs WT.

**Suppl Figure 3.**
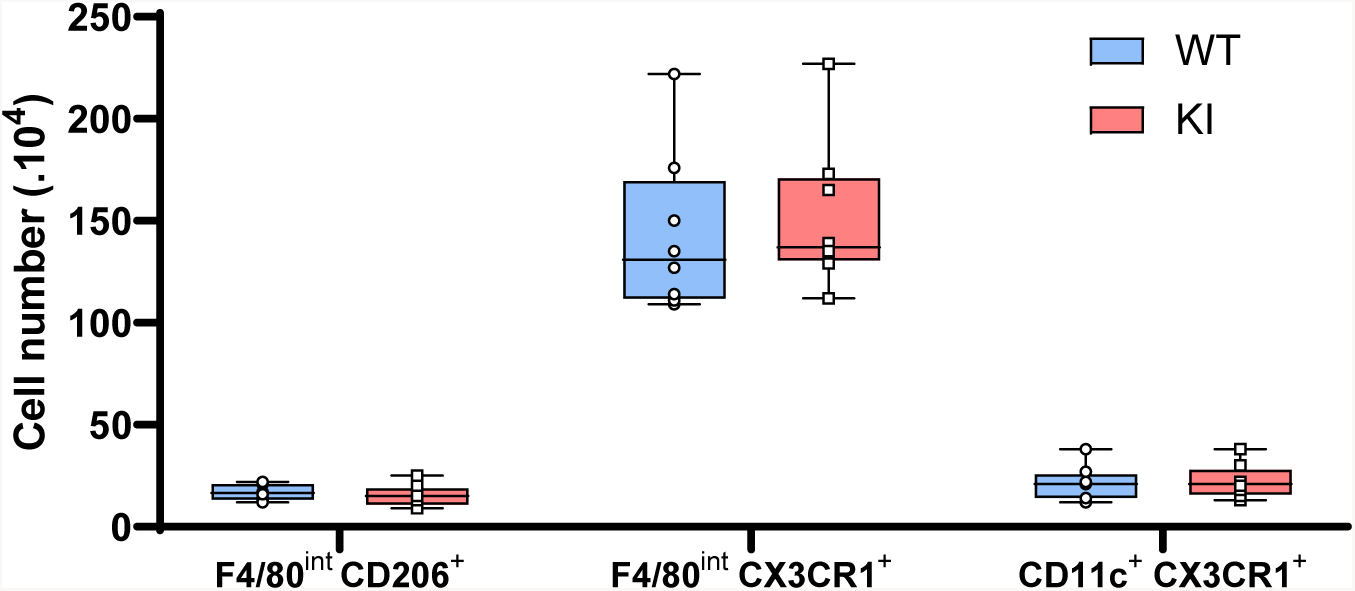
Flow cytometry analysis of macrophage/dendritic cell subsets in the spleens from SOCS2^KI^ or WT mice. Flow cytometry analysis of macrophage/dendritic cell subsets in the spleens from two-month old male SOCS2^KI^ or WT mice (N=8). Statistical analysis was performed using the multiple t-test of ANOVA.

**Suppl Figure 4.**
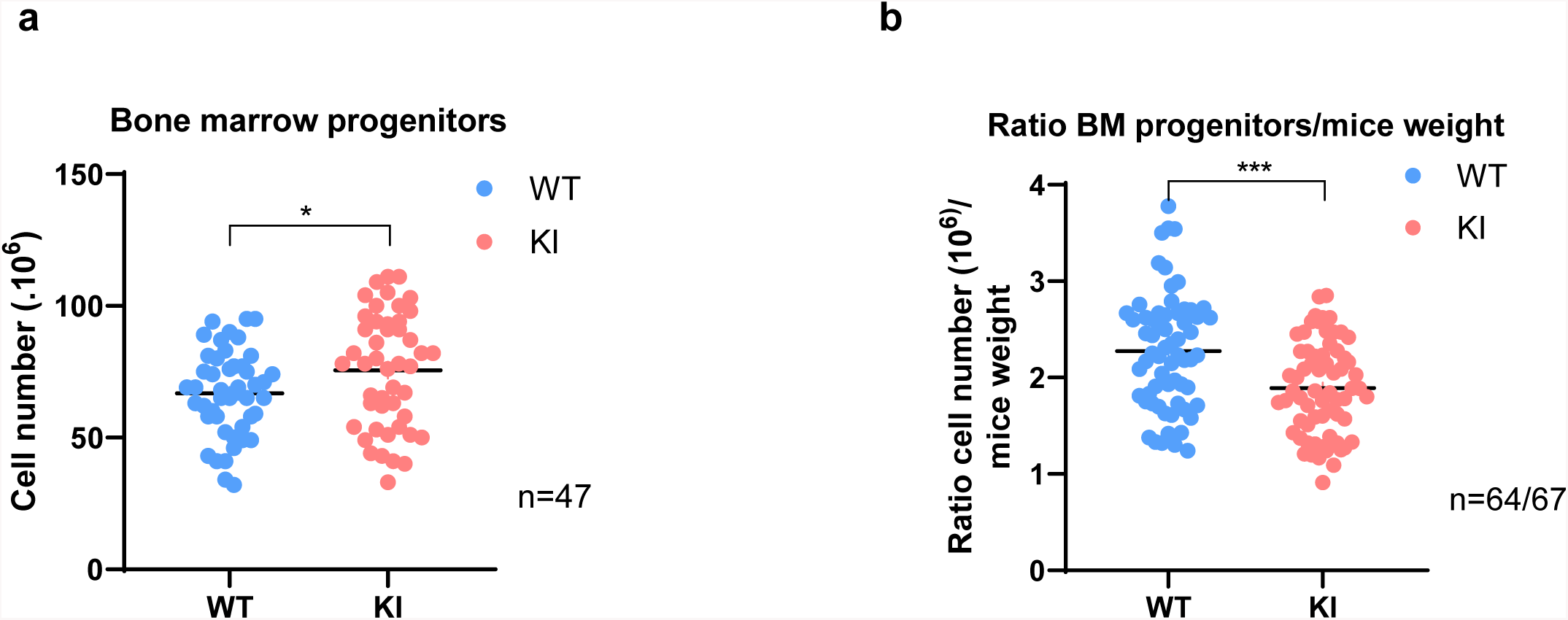
Cell number of bone marrow progenitors from SOCS2^KI^ or WT mice. (a) Cell number of bone marrow progenitors from two-month-old male SOCS2^KI^ or WT mice (N=47). (b) Ratio of the cell number of bone marrow progenitors/mouse weight of SOCS2^KI^ or WT mice (N=64/67). Statistical analysis was performed using the t-test and significant p values are indicated. *P<0.05, **P<0.01. ***P<0.001 vs. WT.

**Suppl Figure 5.**
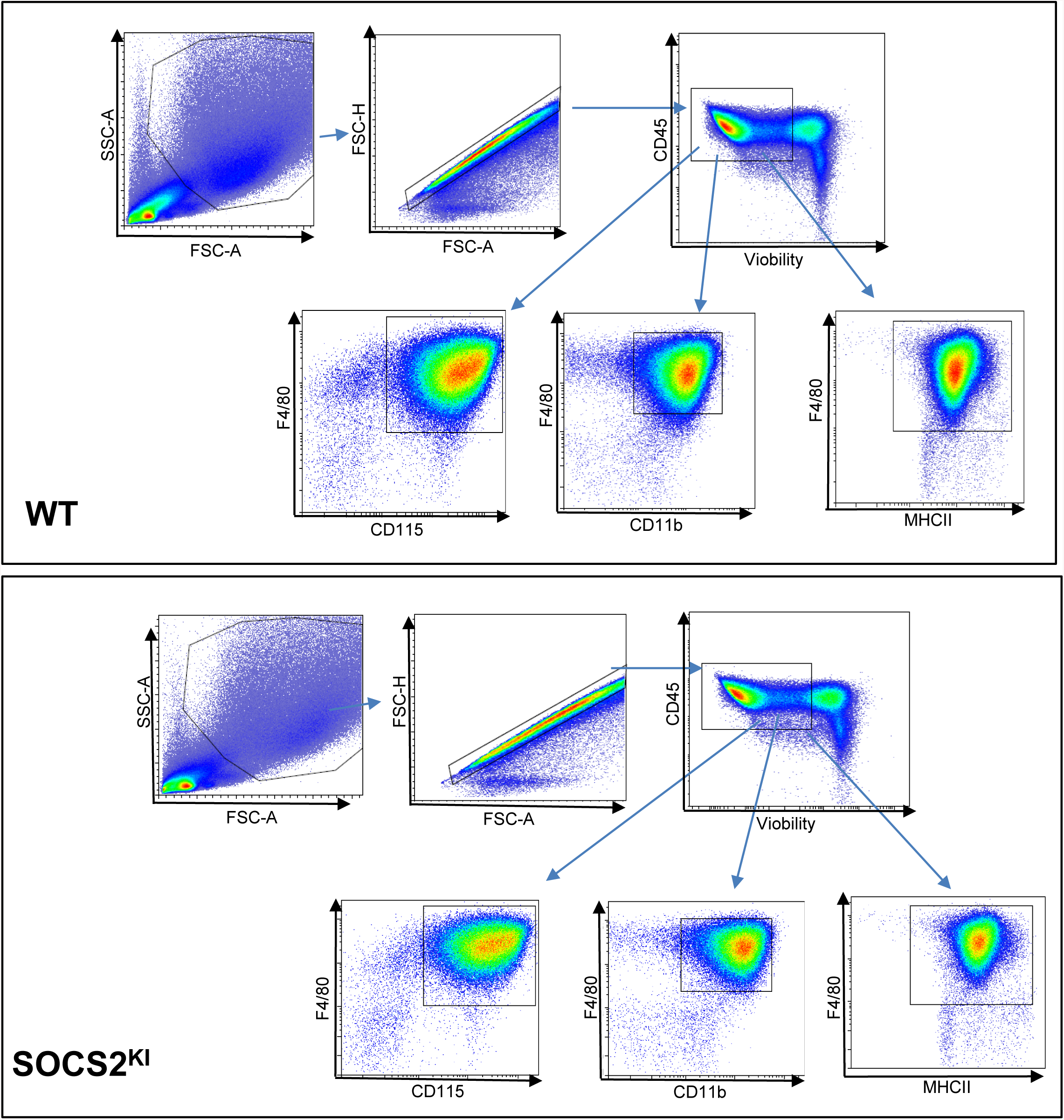
Phenotypic analysis of M-CSF derived BMMs from SOCS2^KI^ or WT mice. Bone marrow progenitors from SOCS2^KI^ and WT mice were cultured with 10 ng/ml of M-CSF for seven days. Adherent cells were harvested and analyzed by flow cytometry for F4/80, CD115, CD11b, and CD11c expression. SOCS, suppressor of cytokine signaling; WT, wild type; KI, SOCS2^KI^ mice.

**Suppl Figure 6.**
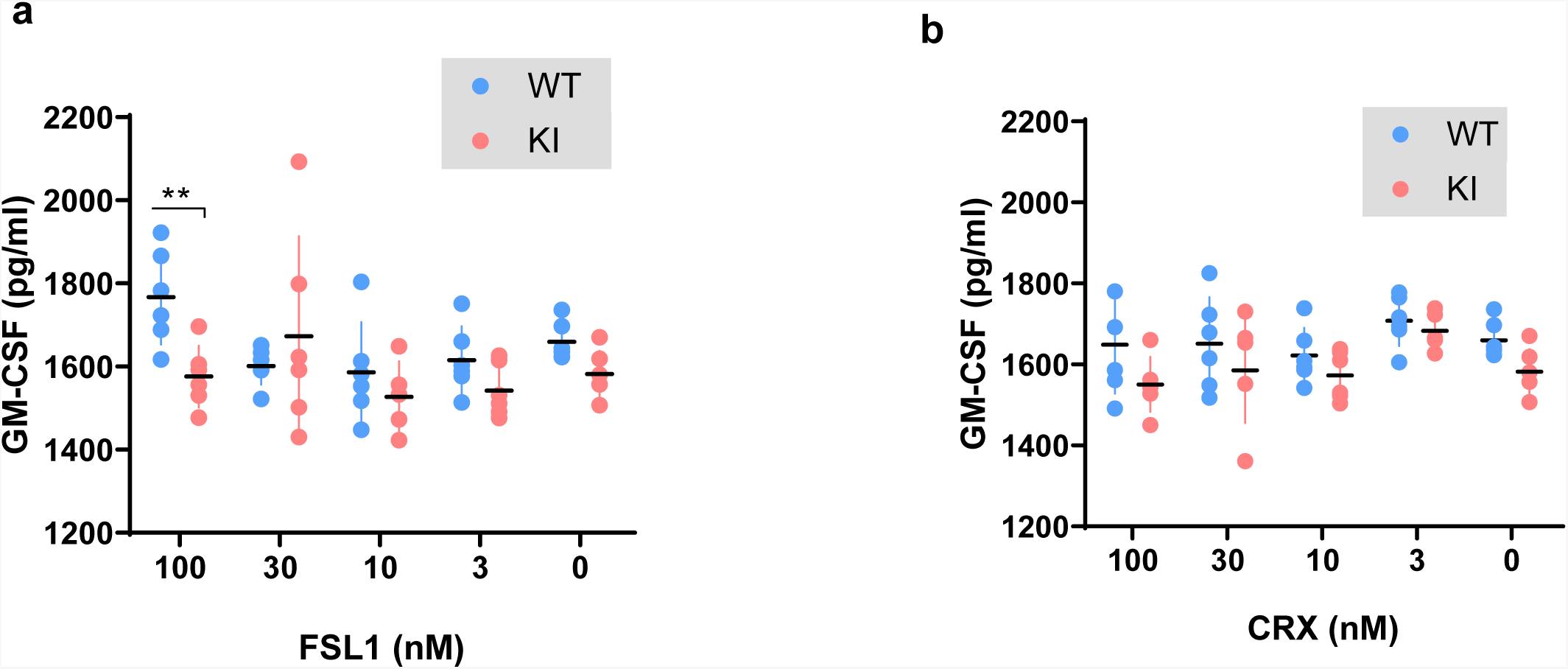
GM-CSF concentrations in BMM supernatant after TLR stimulation. BMMs from SOCS2^KI^ or WT mice were stimulated 24 h with 3 to 100 nM FSL1 or CRX ligands. GM-CSF concentrations were measured in cell supernatants (N=5). Statistical analysis was performed using the multiple t-test of ANOVA. **P<0.01 vs. WT.

**Suppl Figure 7.**
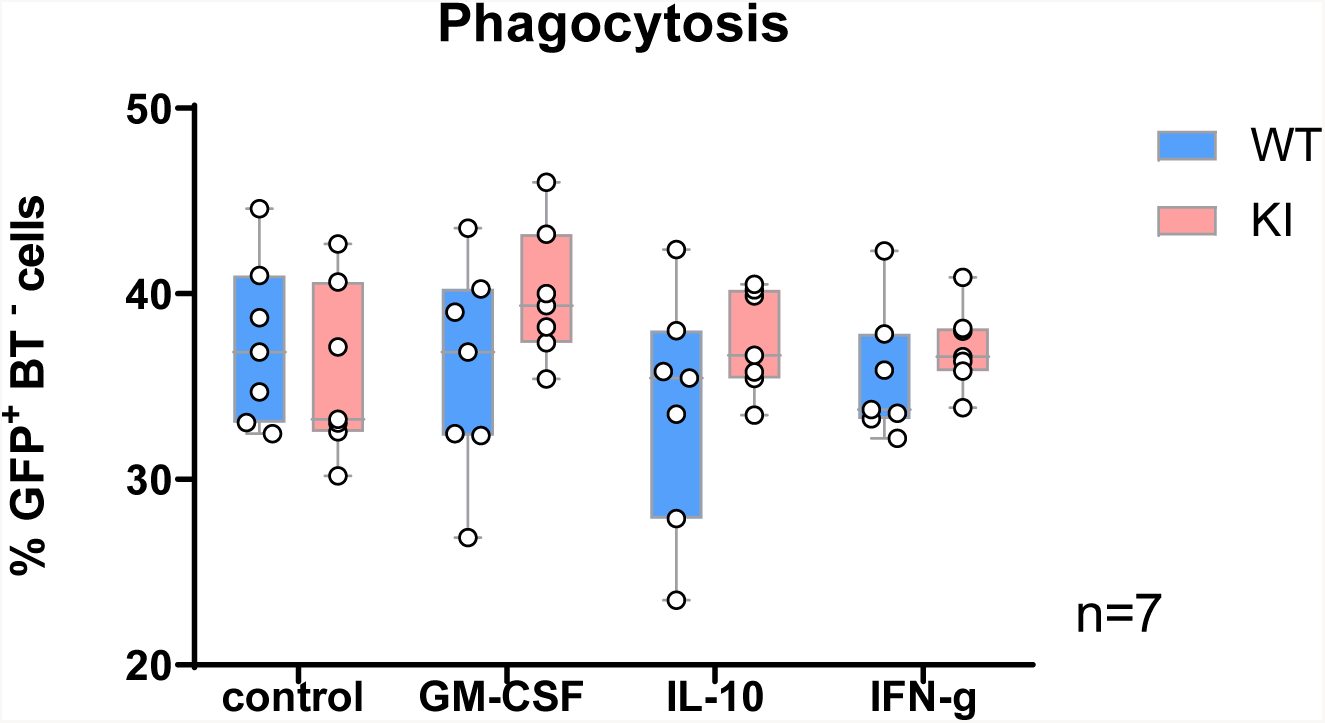
Phagocytic response of SOCS2^KI^ and WT BMMs to the *S. aureus*-GFP strain. BMMs were first primed with either GM-CSF, IFN-γ, or IL-10 for 24 h and then infected with *S. aureus* HG001-GFP for 1 h at 37°C or 4°C as control. The amount of live and dead engulfed bacteria was determined using the GFP^+^7AAD^-^ and GFP^+^ 7AAD^+^ gates, respectively (N=7). Statistical analysis was performed using the multiple t-test of ANOVA and significant p values are indicated. *P<0.05.

**Suppl Figure 8.**
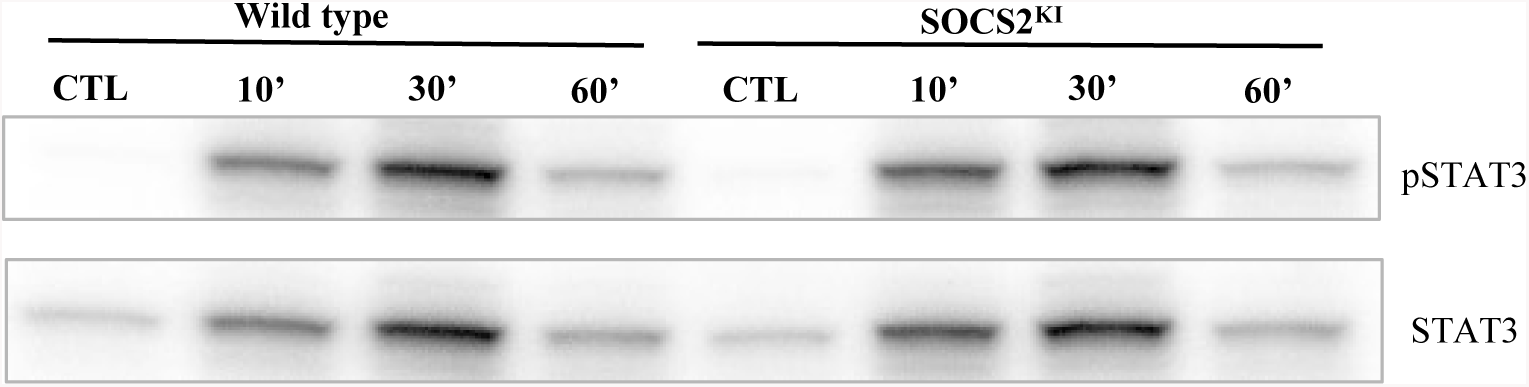
pSTAT3 and STAT3 expression in BMMs from WT vs. SOCS2^KI^ mice after IL-6 stimulation. BMMs were cultured in the presence of GM-CSF (10 ng/ml). The expression of pSTAT3 and total STAT3 were analyzed after stimulation with IL-6 at 20 ng/ml for 10 to 60 min by western blotting. Data are representative of two independent experiments.

**Suppl Figure 9.**
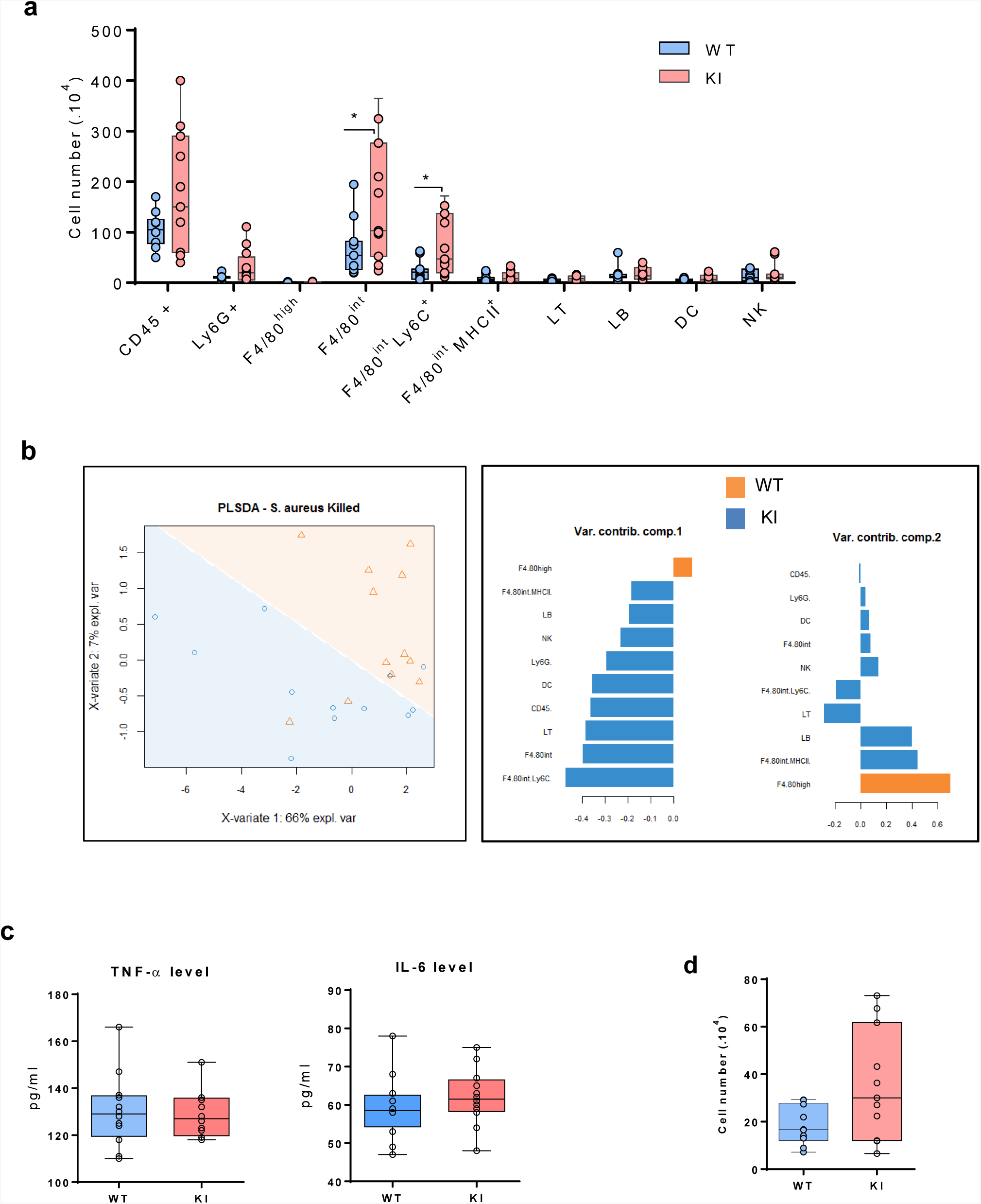
Analysis of the immune response after 16 h of killed *S. aureus* peritonitis in SOCS2^KI^ or WT mice. (**a**) Immune cell-composition in the peritoneal cavity of SOCS2^KI^ or WT mice. (**b**) PSLDA and hierarchical clustering of individual mice as a function of immune cell-composition. The respective contribution of the quantitative variables to dimension 1 and 2 was determined. (**c**) Cytokine concentrations of TNF-α and IL-6 (pg/mL) in exudates. (**d**) Total F4/80^+^ Ly6G^+^ cells /CD45^+^ cells in the peritoneal cavity. Statistical analysis was performed using the t-test and multiple t-test of ANOVA and significant p values are indicated. *P<0.05 vs. WT.

**Supplemental Figure 10 (WB from figure 2).**
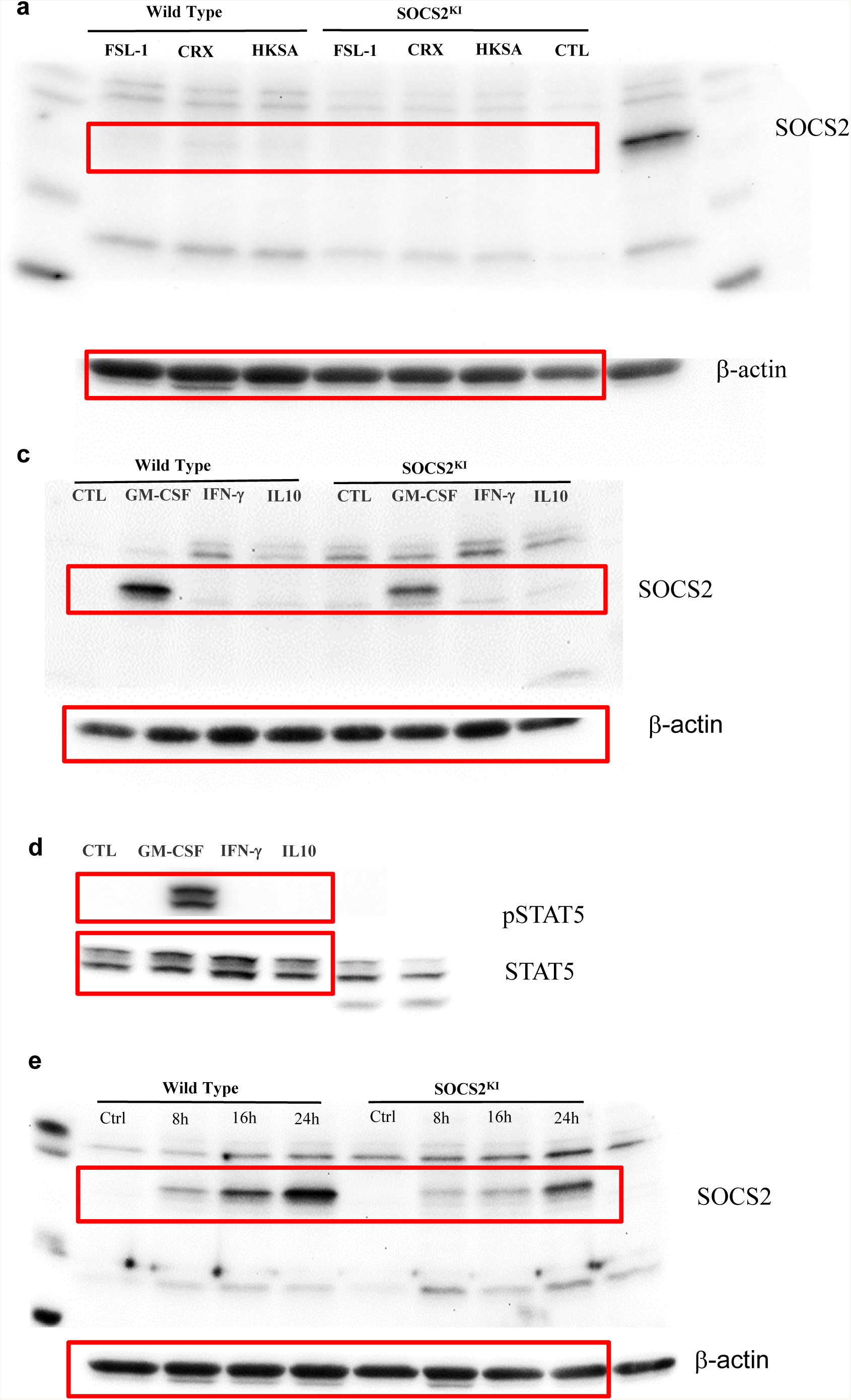

**Supplemental Figure 11 (WB from figure 4).**
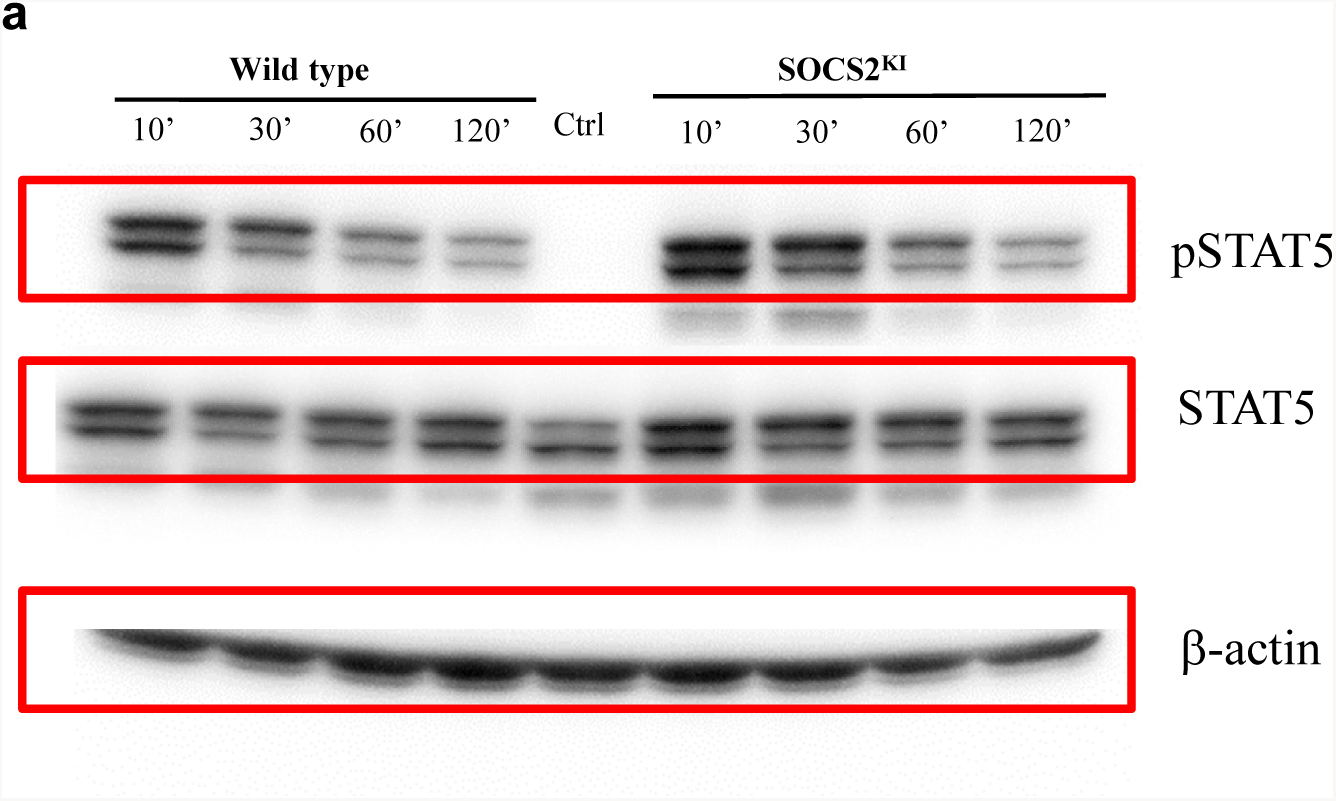

---

**Table 1.** List of antibodies used in the study.

| Antigen | Fluorochrome | Clone | source |
| --- | --- | --- | --- |
| Ly-6G | VioBlue | REAffinity™, clone REA526 | Miltenyi Biotec |
| CD45 | VioGreen | REAffinity™, clone REA737 | Miltenyi Biotec |
| CD335 (NKp46) | PE | REAffinity™, clone REA815 | Miltenyi Biotec |
| CD19 | PE-Vio 615 | REAffinity™, clone REA749 | Miltenyi Biotec |
| CD11c | PE-Vio 770 | REAffinity™, clone REA754 | Miltenyi Biotec |
| F4/80 | APC | REAffinity™, clone REA126 | Miltenyi Biotec |
| CD3 | APC-Vio770 | REAffinity™, clone REA641 | Miltenyi Biotec |
| Ly-6C | VioBlue | REAffinity™, clone REA796 | Miltenyi Biotec |
| Ly-6G | FITC | REAffinity™, clone REA526 | Miltenyi Biotec |
| CD335 (NKp46) | FITC | REAffinity™, clone REA815 | Miltenyi Biotec |
| CD19 | FITC | REAffinity™, clone REA749 | Miltenyi Biotec |
| CD3 | FITC | REAffinity™, clone REA641 | Miltenyi Biotec |
| CD206 (MMR) | PE | Rat, C068C2 | Biolegend |
| CX3CR1 | PerCP/Cyanine5.5 | SA011F11 | Biolegend |
| CD64 | APC-Vio770 | REAffinity™, clone REA286 | Miltenyi Biotec |
| CD11b | APC-Vio770 | REAffinity™, clone REA592 | Miltenyi Biotec |
| MHCII | VioBlue | REAffinity™, clone REA813 | Miltenyi Biotec |

